# Hippocampus consolidates memory in the upstate of cortical sleep slow oscillations

**DOI:** 10.64898/2026.04.17.719155

**Authors:** Maximilian Harkotte, Marion Inostroza, Jan Born, Niels Niethard

## Abstract

Cortical slow oscillations (SOs), a hallmark of non-rapid eye movement (NonREM) sleep, have been proposed to support systems memory consolidation by organizing hippocampal-cortical communication. However, whether consolidation requires hippocampal memory processing during SO-defined windows is unclear. Here, we used closed-loop optogenetics to transiently inhibit dorsal hippocampal activity in adult rats (N = 12) during NonREM sleep following object-place association learning, either during cortical SO upstates or outside SOs, compared with a no-stimulation control. Inhibition during SO upstates completely abolished expression of memory at retrieval, despite preserved sleep architecture and intact cortical SO and spindle dynamics. By contrast, inhibition outside SOs preserved memory and only slightly reduced performance compared to the no-stimulation control. Memory impairment from hippocampal inhibition was largely mediated by SO upstates nesting spindles. Our findings provide novel evidence that sleep-dependent systems consolidation requires precisely timed hippocampal-neocortical dialogue.

## Main

Sleep supports the consolidation of newly acquired memories ^1,2^. Consolidation during sleep is commonly thought of as a systems consolidation process based on the dialogue between the hippocampus and neocortical networks during non-rapid eye movement (NonREM) sleep ^3–6^. Within this framework, cortical slow oscillations (SOs) are assumed to favor the neuronal replay of newly encoded memory representations in hippocampal networks, thereby facilitating the transfer of hippocampally reactivated information as well as memory storage in neocortical networks ^7–9^. SOs emerge in the neocortex as prominent, discrete events consisting of a hyperpolarized downstate followed by a depolarized upstate, recurring at approximately 0.1-4 Hz ^10,11^. At the behavioral level, SOs have been consistently linked to enhanced memory performance ^2,12,13^. At the neural level, the depolarizing SO upstate defines temporal windows during which activity across widespread neuronal populations becomes synchronized, likely driven by the highly coordinated neuronal silence during the preceding downstate ^14,15^. Accordingly, SOs are assumed to also provide periods of enhanced coordination between hippocampal and neocortical neural activity ^16–18^. Indeed, within hippocampal networks, neurons encoding recent experiences are repeatedly reactivated in conjunction with sharp-wave ripples which preferentially occur in close proximity to the SO downstate ^19–21^. Recent work has identified a subset of high-amplitude, long-duration sharp wave-ripples that preferentially emerge during the SO upstate and are associated with enhanced memory reactivation in both the hippocampus and prefrontal cortex ^22^. The SO upstate also promotes the generation of thalamo-cortical sleep spindles ^11,23,24^, which facilitate sharp wave-ripple occurrence during the upstate ^6,25^ and are themselves critical for memory consolidation ^26,27^.

Whereas the role of a hippocampal-cortical dialogue in memory consolidation has been inferred based on correlative findings, direct causal demonstrations remain limited. In a seminal study, Maingret et al. ^28^ showed that electrical stimulation to the neocortex timed to hippocampal sharp wave-ripples enhances hippocampal-cortical coupling as well as later memory retrieval in rats. Here, we provide complementary novel evidence for a causal role of the hippocampal-neocortical dialogue in memory consolidation by adopting the converse approach, i.e., by optogenetically silencing hippocampal activity timed to online detected neocortical SO upstates. Object-location memories were preserved at retrieval 3 hours after encoding when the hippocampus was silenced during post-encoding NonREM sleep outside SOs or in a no-stimulation control condition. In contrast, hippocampal silencing during SO upstates entirely abolished the expression of object-location memory. Mediation analysis indicates that the memory deficit is largely explained by disruption of spindles nesting within SO upstates, implicating SO-spindle-coupled hippocampal processing as the principal mechanism organizing memory consolidation.

## Results

Experiments were performed in male rats (N = 12) which were chronically implanted with EEG screw electrodes (of which the 0.1-4 Hz filtered EEG over left frontal cortex was used for online detection of SO events) and optic fibers positioned bilaterally above the dorsal CA1 (expressing the red-shifted light-activated chloride pump Jaws) for optogenetic inhibition of hippocampal activity (Fig. 1a and Extended Data Fig. 1a). Animals were repeatedly subjected to a standard object-place recognition (OPR) task consisting of a 10-min encoding phase, a 3-hour retention interval during which the animals slept in a resting box, and a 5-min retrieval phase (Fig. 1b). Closed-loop optogenetic inhibition of the hippocampus was delivered either during the SO upstate (In-Phase condition), outside periods of online-detected SOs (Out-of-Phase condition), or not delivered despite SO detection (No-Stimulation control condition, Fig. 1c), using a within-subject-design (Extended Data Fig. 1b). The efficacy of hippocampal inhibition by Jaws activation was tested in three additional animals and confirmed that light delivery acutely produced a robust and consistent suppression of neuronal firing rates (Wilcoxon signed-rank test: *V* = 6486.5, *p* < 0.001), with 84.5% of recorded neurons showing suppressed activity during illumination (Fig. 1d,e, Extended Data Fig 1c,d).

**Fig. 1.**
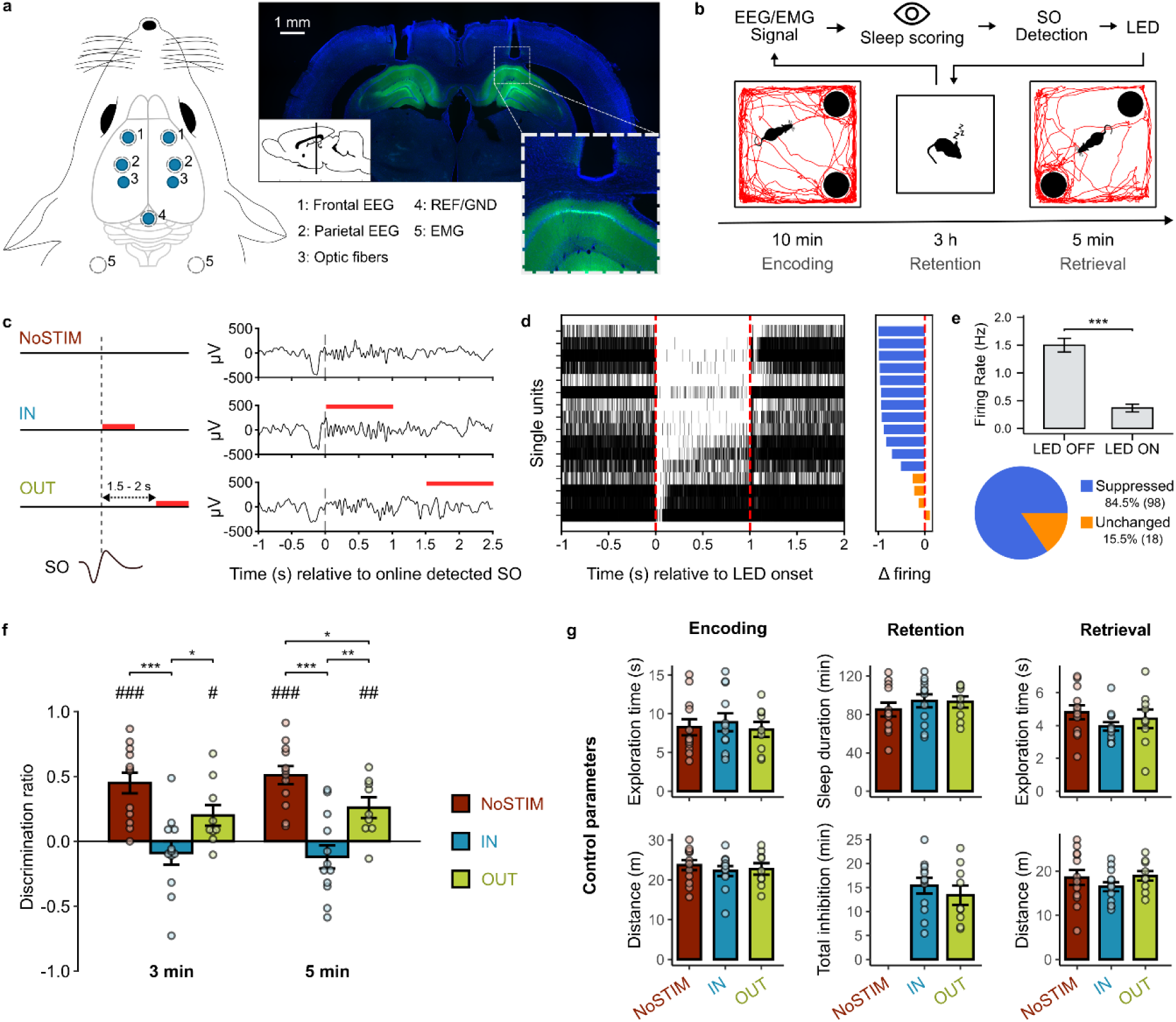
Timing of hippocampal inhibition during cortical SOs determines its impact on memory consolidation. a, Left, Chronic implantation of frontal, parietal and occipital EEG screw electrodes (reference/ground), bilateral optic fibers above dorsal hippocampus, and neck EMG electrodes. Right, Bilateral optic fiber placement above dorsal CA1 expressing the red-light-activated chloride pump Jaws-GFP. **b,** Object-place recognition task. During encoding, rats explored two identical objects for 10 min. During the 3 h retention interval, cortical SOs were detected online during NonREM sleep to trigger LED illumination. During the 5 min retrieval test, one object was displaced to a novel location. **c,** Closed-loop hippocampal inhibition during retention: No-Stimulation control (NoSTIM, top), inhibition during the SO upstate (IN, middle), or during NonREM sleep outside of SOs (OUT, bottom). Representative EEG traces illustrate LED timing (red bars). **d,** Left, Suppression of hippocampal spiking in one rat under urethane anesthesia (spikes of 16 single units summed across 593 trials). Right, Normalized spike-count differences between inhibition and baseline windows (units sorted by suppression magnitude). **e,** Top, Firing rates of 116 single units (N = 3 rats) recorded under urethane anesthesia decrease during LED illumination. Bottom, Proportion of suppressed units and unchanged units. **f,** Mean ± SEM cumulative discrimination ratios during retrieval shows better spatial memory in the No-Stimulation control condition than in the Out-of-Phase-inhibition condition, whereas inhibition during the SO upstate abolishes memory performance (*p< 0.05, **p < 0.01, ***p< 0.001 for condition comparisons; ^##^p< 0.01, ^###^p< 0.001 against chance). **g,** Control measures: total object exploration time and distance traveled during encoding and retrieval (left, right), as well as total sleep duration and total inhibition time during retention (middle), were comparable across conditions.

### Hippocampal inhibition during cortical SO upstates disrupts OPR memory consolidation

Memory performance during the OPR retrieval phase was quantified using the discrimination ratio, reflecting the relative exploration time of the object in the novel versus familiar location. Replicating previous reports ^29–32^ animals exhibited robust memory performance in the No-Stimulation control condition, as indicated by a high discrimination ratio during the first three and full five minutes of the test phase (mean ± SEM: 0.45 ± 0.08 and 0.51 ± 0.07, respectively; *t*(11) = 5.33 and *t*(11) = 7.07, *p* < 0.001 against chance level; Fig. 1f). In contrast, optogenetically inhibiting the hippocampus In-Phase, i.e., during the SO upstates of post-encoding NonREM sleep, effectively abolished memory performance (mean ± SEM: −0.09 ± 0.09 and −0.12 ± 0.09, respectively for three and five minutes; *t*(11) = −1.04, *p* = 0.32 and *t*(11) = −1.25, *p* = 0.24 against chance level; *t*(20.00) = 4.44, *p* < 0.001 and *t*(20.60) = 5.29, *p* < 0.001 for post-hoc comparison against the No-Stimulation condition). Notably, memory performance in the Out-of-Phase inhibition condition remained significantly above chance (mean ± SEM: 0.20 ± 0.08 and 0.26 ± 0.08, respectively for three and five minutes; *t*(8) = 2.44, *p* = 0.04 and *t*(8) = 3.42, *p* = 0.009 against chance level), and also was significantly higher than in the In-Phase condition (*t*(18.80) = 2.44, *p* = 0.05 and *t*(18.90) = 3.12, *p* = 0.006, respectively for three and five minutes), but modestly reduced compared to the No-Stimulation condition (*t*(18.60) = −2.06, *p* = 0.054 and *t*(18.20) = −2.40, *p* = 0.03, respectively for three and five minutes). The overall effect of inhibition condition was additionally confirmed by an analysis that controlled for the order of stimulation conditions (χ²(2) = 45.06, *p* < 0.001; Extended Data Fig. 1e). Importantly, we did not find any differences between conditions in control measures, including distance travelled and total object exploration time during encoding and retrieval, as well as total sleep duration, total inhibition time, mean inhibition time, and inhibition density during the retention interval (all *p* > 0.05, Fig. 1g and Extended Fig. 1f), thereby excluding confounding effects of nonspecific arousal-related factors on memory performance. These results indicate that the timing of optogenetic hippocampal inhibition during NonREM sleep critically determines its impact on memory consolidation, with the SO upstate representing the most sensitive window.

### Macro sleep architecture as well as cortical SO and spindle dynamics are preserved despite hippocampal inhibition

Memory consolidation has been shown to correlate with total sleep time and the amount of NonREM sleep ^33,34^, and the magnitude of EEG slow-wave activity during NonREM sleep ^35^, suggesting that both sleep duration and depth contribute to consolidation. Accordingly, we examined whether closed-loop hippocampal inhibition altered overall sleep architecture during the post-encoding interval. Across the 3-hour retention period, animals spent approximately 88.34 ± 21.17 min awake, 81.77 ± 18.92 min in NonREM sleep and 6.82 ± 3.96 min in REM sleep (mean ± SD), with no differences between conditions (all *p* > 0.05, Fig. 2b). Likewise, the average duration of individual Wake, NonREM, and REM epochs (mean ± SD: 94.78 ± 36.81, 86.48 ± 24.62, and 118.52 ± 34.66 s) as well as latencies to NonREM and REM sleep onset (mean ± SD: 19.64 ± 18.92 and 91.70 ± 3.96 min, respectively) were comparable across conditions (all *p* > 0.05, Extended Data Fig. 2a). Spectral power distributions during NonREM and REM sleep (from left frontal EEG) were indistinguishable between conditions as well (all p > 0.05, Fig. 2c).

**Fig. 2.**
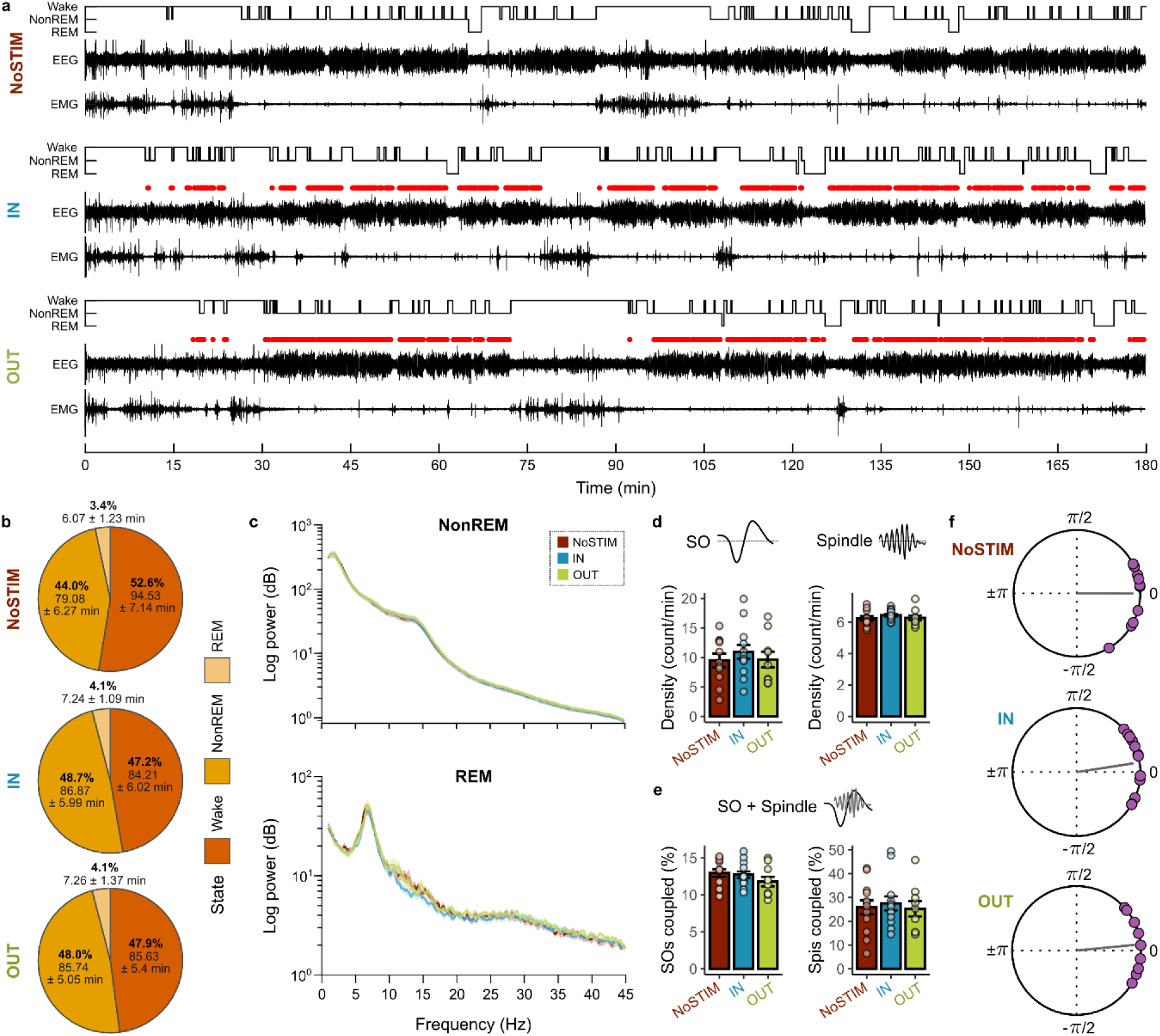
Preserved macro sleep architecture and cortical oscillatory dynamics despite hippocampal inhibition. a, Representative hypnograms of one animal during the 3 h retention interval with raw frontal EEG, EMG, and optogenetic inhibition times (red dots). Top, No-Stimulation control condition (NoSTIM); Middle, In-Phase inhibition (IN); Bottom, Out-of-Phase inhibition (OUT). EEG scale: ±500 µV; EMG scale: ±1000 µV. **b,** Mean ± SEM total time spent in each brain state during retention were comparable across conditions (condition comparison: Wake: χ^2^(2) = 4.99, *p* = 0.08, NonREM: χ^2^ (2) = 3.47, *p* = 0.18, REM: χ^2^(2) = 1.91, *p* = 0.39). **c,** Mean spectral power during NonREM (top) and REM (bottom) sleep epochs did not differ between conditions. **d,** Mean density of online-detected cortical SOs (left) and sleep spindles (right) during the retention interval were comparable across conditions. **e,** SO-spindle coupling, defined as spindle onsets occurring within 1 s after online SO detection, was comparable across conditions, both for the percentage of SOs followed by spindles (left) and the percentage of spindles coupled to SOs (right). **f,** Mean SO phase at which spindle power peaked. Purple dots indicate individual animals; grey lines indicate the mean phase of maximal spindle power. In all conditions, spindle power peaked near the SO positive maximum (0°, Watson-Williams Test for condition comparison: *p* = 0.74).

Considering evidence that hippocampal activity, particularly the occurrence of sharp wave-ripples, may influence the emergence of cortical SOs in a bottom-up manner ^26,36,37^, we examined whether closed-loop optogenetic inhibition of the dorsal CA1 altered cortical SOs, spindles, or their temporal coupling. As sharp-wave ripples and associated hippocampal memory replay occurs time locked to SOs ^20,38^, stronger changes in SO and spindle dynamics might have been expected for the In-Phase inhibition of CA1 compared to the Out-of-Phase condition. However, the three experimental conditions did neither differ in SO density or negative-to-positive peak amplitude, nor in spindle density or absolute peak amplitude (all p > 0.05, Fig. 2d and Extended Data Fig. 2b). Also, SO-spindle coupling remained unaffected: The percentage of SOs coupled to spindles, defined by a spindle peak occurring within 1 s following SO detection, was comparable across conditions, as were the percentage of coupled spindles (all p > 0.05, Fig. 2e). Likewise, phase-amplitude coupling, i.e., the exact SO phase at which spindle peaks occurred, was close to the SO positive maximum and comparable in all three conditions (Watson-Williams Test, *p* = 0.74, Fig. 2f). These results for the frontal EEG were replicated for parietal EEG channels located in closer proximity to the hippocampus (Extended Data Fig. 2c-e). Together, these findings demonstrate that closed-loop hippocampal inhibition - whether applied during SO upstates or outside of SOs - did not substantially alter cortical SO or spindle dynamics, or any relevant parameters of sleep macro-architecture.

### Memory impairment following closed-loop hippocampal inhibition during SO upstates is mediated by cortical spindles

SOs often nest a spindle in their upstate and there is some evidence that, rather than directly impacting hippocampal memory processing, the influence of SO upstates may be conveyed via thalamically generated spindles ^6,11,36^. For example, optogenetic induction of spindles temporally grouped hippocampal ripples, independent of the SO phase in which they were induced ^39^. Indeed, time-frequency plots aligned to the rising flank of online-detected SOs revealed a robust increase in spindle-frequency power (10-16 Hz) during the SO upstate in the No-Stimulation control condition (Extended Data Fig. 3a) as well as in the In-Phase condition, in which the rising SO flank was equivalent to the inhibition onset, whereas in the Out-of-Phase condition, this increase in spindle power preceded the inhibition window (all clusters p < 0.05; Fig. 3a). To examine whether the impairment of OPR memory following hippocampal inhibition during the SO upstates reflected a suppression of spindle-mediated, rather than direct influences of cortical SOs on hippocampal networks, we performed a multilevel mediation analysis. This analysis tested whether the difference in retrieval performance between the In-Phase and Out-of-Phase conditions can be explained by the percentage of spindles that overlapped with the inhibition window. Indeed, the two conditions differed in the percentage of spindles affected by hippocampal inhibition (*β* = 0.14 ± 0.03, *t* = 5.48, *p* < 0.001, Fig. 3b,c), and a higher fraction of inhibited spindles predicted poorer retrieval performance (*β* = −2.20 ± 0.85, *t* = −2.59, *p* = 0.02). The mediation analysis revealed a significant indirect, spindle-mediated effect (average causal mediation effect, ACME = −0.30, 95% CI −0.57 to −0.07, *p* = 0.008), which accounted for approximately 84% of the total effect (*p* = 0.01). In contrast, the direct effect of inhibition condition after accounting for spindle disruption was not significant (*β* = −0.07 ± 0.15, *t* = −0.47, *p* = 0.64). Consistent effects were also observed in parietal EEG (Extended Data Fig. 3b-d). Although these results support the view that spindles in the SO upstate mediate the effect of In-Phase inhibition of hippocampal CA1 on memory consolidation, they do not rule out that spindles themselves, i.e., in absence of SOs are able to trigger consolidation of OPR memory. However, memory performance did neither correlate with the number of solitary spindles, i.e., spindles occurring in the absence of a SO, in any of the three experimental conditions (all *R* < 0.29, *p* > 0.39, uncorrected for multiple comparisons), nor with the fraction of spindles occurring during the inhibition window in the Out-of-Phase condition (*R* = −0.15, *p* = 0.71). Taken together, these findings point to hippocampal-cortical interactions supporting OPR memory consolidation being primarily mediated by hippocampal processing during SO-spindle events.

**Fig. 3.**
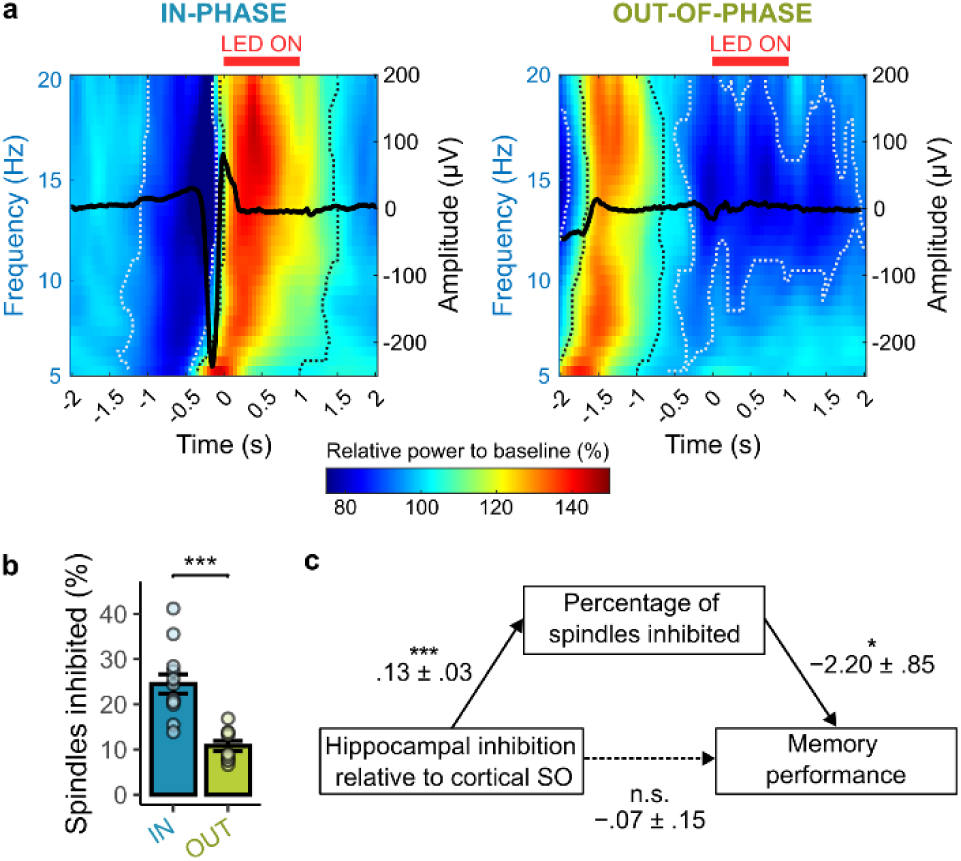
Memory impairment induced by hippocampal inhibition during SO upstates is spindle-mediated. a, Time-frequency representations locked to the inhibition onset in the In-Phase (left) and Out-of-Phase (right) conditions. Dashed lines indicate significant positive (black) and negative (white) spectral clusters relative to baseline. **b,** The fraction of all spindles overlapping with the hippocampal inhibition window was significantly higher in the In-Phase than in the Out-of-Phase condition (*t*(16.30) = 5.34, *p* = 0.054, ***p< 0.001). **c,** Mediation model relating inhibition condition (In-Phase versus Out-of-Phase), the percentage of inhibited spindles, and memory performance. Path coefficients represent unstandardized fixed-effect estimates (estimate ± s.e.) from linear mixed-effects models with animals included as a random intercept. Solid arrows indicate significant paths; the dashed arrow denotes the direct effect of condition after accounting for the mediator (*p< 0.05, ***p< 0.001).

## Discussion

According to active systems consolidation theory, memory consolidation during sleep relies on a coordinated dialogue between neocortical and hippocampal networks, with neocortical slow oscillations (SOs) providing temporal windows for hippocampal reactivation and information transfer to long-term cortical stores ^2,4^. While there is a large body of evidence indicating causal contributions to memory consolidation of both neocortical SOs ^12,40,41^ as well as hippocampal memory reactivations ^42–44^, evidence for the causal importance of a hippocampal-neocortical dialogue mediating consolidation during sleep remains scarce ^28^. Here, we provide such evidence, showing that effective consolidation of spatial object-location memory requires hippocampal activity during SO upstates. Inhibiting dorsal CA1 during SO upstates (In-Phase inhibition) in post-encoding NonREM sleep completely abolished behavioral expression of object-location memory at a later retrieval test. With inhibitions of CA1 outside of SOs (Out-of-Phase inhibition) object-place memory was preserved, although slightly diminished in comparison with the No-Stimulation control condition. The mediation analysis further suggests that the effect of SOs on hippocampus-dependent consolidation of memories is primarily conveyed through spindles nesting into the SO upstate. The findings extend prior work by providing direct causal evidence that temporally precise hippocampal activity during the SO upstate is required for sleep-dependent memory consolidation.

The impairing effect on spatial memory consolidation was specific to hippocampal inhibition during SO upstates. Brief optogenetic inhibition of CA1 neuronal activity was triggered in real time by closed-loop detection of cortical SOs during post-encoding sleep. The 1-s inhibition window was chosen to encompass the entire SO upstate including the period of maximal spindle power. Although inhibition during this window abolished memory consolidation, it altered neither sleep macro-architecture nor the incidence or coupling of SOs and spindles, ruling out that the observed memory impairment was due to nonspecific confounds of inhibition.

SOs originating from neocortical networks can travel to the hippocampus to directly impact memory processing in these networks ^45–48^. However, neocortical SOs may also indirectly affect hippocampal memory processing through spindles nesting in the SO upstate ^6,36,39^. Our mediation analysis indicates that the memory impairment produced by hippocampal inhibition during SO upstates was primarily linked to disruption of coupled SO-spindle events. Importantly, this mediation does not imply that spindles act independently of SOs, but rather that they constitute a key intermediary mechanism through which SO-defined windows shape hippocampal processing. In line with this, spindle activity occurring outside SOs did not predict behavioral memory performance at retrieval, nor did spindle activity present during periods of hippocampal inhibition in the Out-of-Phase condition. This pattern aligns with previous work in which thalamic spindles were optogenetically induced to examine their role in memory consolidation ^39^. In those experiments, spindles were the primary factor timing hippocampal ripples associated with memory replay, regardless of whether they were induced during an online-detected SO upstate or outside any SO. Notably, however, intact spatial memory at later retrieval required that spindle induction coincided precisely with the SO upstate, whereas spindles induced outside SOs did not enhance memory. Together, these findings support the view that the temporally organizing influence of SOs on hippocampal memory reactivations is primarily conveyed through coupled thalamic spindles. Effective hippocampo-to-neocortical transfer of reactivated memory information and its long-term storage into neocortical circuits, however, require that spindle-timed hippocampal memory reactivations occur during the excitable upstates of neocortical SOs ^49^.

Inhibiting hippocampal activity during NonREM sleep outside of SOs slightly reduced memory performance, but retrieval remained above chance. In this Out-of-Phase condition, hippocampal inhibition was delayed (by 1-2 s) relative to SO detection and terminated upon the occurrence of a subsequent SO or transition into wake or REM sleep. Hippocampal inhibition in this condition closely matched with that in the experimental In-Phase condition differing only in the timing of the inhibition relative to cortical SOs (Fig. 1g and Extended Data Fig. 1f). The slight decrease in memory in this Out-of-Phase condition was unexpected, but might hint at hippocampal processing outside of SOs that adds to memory stabilization. For example, spindles might be implicated in memory consolidation independent of co-occurring SOs ^16,27,50,51^, although our correlational analyses did not reveal any significant link between spindle activity during Out-of-Phase hippocampal inhibition and memory performance. Alternatively, the optogenetic inhibition may have affected intrahippocampal processes that support memory consolidation independently of both SOs and spindles. Recent work, for example, indicates that memory reactivation during post-learning NonREM sleep is also regulated by interneuron-mediated mechanisms within the hippocampus that operate outside periods of sharp-wave ripples and help balance replay-related activity ^52^. Finally, the memory decrease in the Out-of-Phase condition could also reflect a limitation of our closed-loop approach, which detected SOs only when exceeding a certain amplitude criterion and, thus, might have missed smaller, more local SOs that nevertheless could impact hippocampal memory processing ^53,54^.

In sum, our findings identify the SO upstate as a critical temporal window of hippocampal-cortical interaction underlying effective memory consolidation during sleep. At the same time, they point to contributions of hippocampal activity outside of SOs. To what extent these contributions likewise reflect hippocampal-cortical interactions warrants further study.

## Methods

### Animals

Nineteen adult male Long-Evans rats (Janvier, Le Genest-Saint-Isle, France), aged 9-12 weeks at the start of the experiment, were used in this study. The rats were housed in groups of 2-4 per cage, with ad libitum access to food and water throughout the experiment. They were maintained on a 12-hour light/dark cycle (lights on at 6:00 am). Prior to the experiment, the animals were handled daily for 10-15 minutes over five consecutive days. All experimental procedures were conducted in accordance with European animal protection laws and policies and were approved by the Baden-Württemberg state authorities.

### Surgical procedures

Animals underwent two surgical procedures: (i) virus injection four weeks prior to behavioral testing, and (ii) implantation of EEG/EMG electrodes and optic fibers one week before testing, or, in three animals, an acute recording procedure (see *Acute Recordings*).

Before each surgery, rats received an intraperitoneal injection of anesthetic mixture (0.005 mg/kg fentanyl, 2 mg/kg midazolam, and 0.15 mg/kg medetomidine). Surgeries were performed under general isoflurane anesthesia (induction: 1-2%; maintenance: 0.8-1.2% in 0.35 L/min O₂). Rats were placed in a stereotaxic frame, body temperature was maintained at 33-36 °C with a feedback-controlled heating pad, and eyes were protected with ophthalmic ointment, before the skull was exposed.

During the first surgery, animals were bilaterally injected with 500 nL of viral vector (AAV5-hSyn-Jaws-KGC-GFP-ER2; Addgene Plasmid #65014; diluted 1:4 in sterile PBS; resulting titer: ∼1.75 x 10^12^ vg/mL) targeting the dorsal CA1 (AP: −3.8 mm, ML: ±2.4 mm, DV: −2.3 mm, relative to Bregma), to enable AAV-mediated expression of Jaws, a red-shifted light-activated chloride inward pump. Expression was driven by the human synapsin promoter, which supports robust transgene expression across neuronal populations but does not distinguish between excitatory and inhibitory neurons. The virus was delivered at 0.1 µL/min via a sharpened glass pipette (Wiretrol II, Drummond Scientific; tip diameter <25 µm), which remained in place for 10 min post-injection to prevent backflow, then was automatically withdrawn at 0.2 µm/s using a motorized micromanipulator (MP-285, Sutter Instruments). Craniotomies were sealed with silicone elastomer (Kwik-Cast, World Precision Instruments) and cold-polymerizing dental resin (Palapress, Kulzer), and the wound was sutured.

For the second surgery, the sealant and dental resin were removed after skull exposure. Five stainless-steel screw electrodes (Plastics One) were implanted: two frontal (AP: +2.6 mm, ML: ±1.5 mm), two parietal (AP: −1.5 mm, ML: ±2.5 mm), and one occipital (AP: −10.0 mm, ML: 0 mm), the latter serving as reference and ground. Two 400-µm diameter optic fibers (NA 0.5; CFML15L10, Thorlabs) mounted in guide cannulae (OGL, Thorlabs) were bilaterally implanted above dorsal CA1 (AP: −3.8 mm, ML: ±2.4 mm, DV: −1.8 mm). Two stainless-steel wire electrodes were inserted bilaterally into the neck muscles for EMG recording. All electrodes were connected to a Mill-Max pedestal (Mill-Max Mfg. Corp.) and secured to the skull with dental resin. After each surgery, rats received subcutaneous carprofen (5 mg/kg) and were allowed to recover for at least 7 days before the start of behavioral testing.

### Acute recordings

To validate optogenetic inhibition, three animals underwent acute urethane-anesthetized recordings 4-6 weeks after viral injection. A two-shank, 32-channel sharpened silicon probe (P2-ASSY-116, Cambridge Neurotech) was mounted on a custom adapter coupled to a stepper motor actuator (ZST225B, Thorlabs) equipped with a 400-µm diameter optical fiber (NA 0.5; Thorlabs), coupled with a 625 nm LED (M625F2, Thorlabs). This configuration allowed the optical fiber to be positioned <200 µm from the probe shanks and to be moved independently using Kinesis software (Thorlabs).

Anesthesia was induced by stepwise intraperitoneal injection of urethane (1.5 g urethane/5 ml sterile saline; 0.005 ml per g body weight) until no reflexes were observed. Animals then received subcutaneous carprofen (5 mg/kg) and were secured in a stereotaxic frame. The skull was exposed, and sealant and dental resin were removed above the left hemisphere. A screw electrode was implanted at AP = −10.0 mm, ML = 0 mm relative to bregma and connected to the silicon probe to serve as reference and ground. The exposed cortical surface was covered with saline, and the probe-fiber assembly was slowly lowered into the brain under visual guidance to prevent bending of the probe shanks. Once the probe/fiber tips were positioned above CA1 (AP = −3.8 mm, ML = −2.4 mm, DV = −1.8 mm), the assembly was left in place for 1 h to allow the tissue to settle. The probe was then advanced in 50-µm steps at 2 µm/s using the stepper motor until auditory monitoring of the neural activity from the probe indicated proximity to the CA1 pyramidal layer, followed by 20-µm steps until clear spiking activity was detected across all channels. Recordings were then initiated while delivering 0.5-1 s red LED light pulses (10-20 mW at the fiber tip) with randomized 3-5 s inter-trial intervals to avoid rhythmic entrainment of spiking activity and 5-minute breaks after 150 light pulses to let the tissue reset. After a 1 h recording session, the probe-fiber assembly was retracted, and the animal was transcardially perfused (see *Histology*).

### Apparatus and Objects

The object-place recognition (OPR) task was conducted in a square open-field arena (80×80×40 cm) made of gray PVC. The arena was dimly lit (20-30 lux) and supplemented with constant white noise (60 dB). A camera (Logitech C920) was mounted above the arena for video recording. Distal spatial cues included the camera, posters affixed to the walls, hanging objects (e.g., spheres, egg boxes), and surrounding curtains. Adjacent to the arena, a resting box (35×35×45 cm) made of stainless steel and filled with bedding material was used to house the animals during the post-encoding phase. The resting box was enclosed within a Faraday cage, and the animal was monitored using a second camera. Three pairs of glass objects of different shapes and sizes (height: 15-30 cm; base diameter: 7-12 cm), each filled with colored sand, were used. Objects were sufficiently heavy to prevent displacement by the rats. To minimize olfactory cues, both the arena and the objects were thoroughly cleaned with 70% ethanol after each trial.

### Object-place recognition task

Before the experiment, animals were habituated to the experimental context over three consecutive days (Extended Data Fig. 1b). Each day, rats first acclimated to the testing room for 20 min in their home cage before being placed into an empty IVC cage containing an unfamiliar object (distinct from those used in the experiment) for 5 min to familiarize them with the presence of objects. Subsequently, animals were placed for 10 min into the empty open-field arena, facing a different wall on each habituation day. Afterwards, rats were connected to the EEG and optic fiber cables and transferred to the resting box, where they remained undisturbed for 4 h with free access to water (but not food).

For the encoding phases, rats were brought back to the testing room and, after a 20-min acclimation period, placed in the open-field arena containing two identical objects positioned equidistant from two corners. Following 10 min of exploration, the animals were connected to the EEG and optic fiber cables and placed in the resting box for a 3-h retention interval. After this period, rats were disconnected and left undisturbed in the resting box for 5 min to allow grooming before the retrieval phase. During retrieval, the animals were reintroduced into the arena, which contained the same two objects, but one object was relocated to a different corner relative to the encoding session. Animals were subjected to the OPR task either twice (n = 3) or three times (n = 9), each using a different combination of objects and locations within the arena. The first two OPR sessions were separated by 2 days, and the third by 3-7 days, to minimize potential interference between tests.

### Electrophysiological recordings

During behavioral experiments, EEG and EMG signals were acquired via the Mill-Max pedestal and EEG tether (HS-18, Neuralynx) connected to a Digital Lynx SX acquisition system (Neuralynx). Signals were sampled at 1 kHz using Cheetah software (Neuralynx). For acute recordings, a 32-channel silicon probe was connected to the same acquisition system through a buffered 36-channel headstage (HS-36, Neuralynx), and raw data were sampled at 32 kHz.

### Online detection of slow oscillations

Cortical SOs during NonREM sleep were detected online using the left frontal EEG electrode. To determine individualized detection thresholds, EEG recordings from the second habituation session were first sleep-scored (see *Offline sleep scoring*), and SOs were identified offline (see *Offline detection of slow oscillations and sleep spindles*). The mean negative amplitude across offline-detected SOs was then used as an individual negative threshold for online detection during the retention interval (mean ± SD: −120.12 ± 32.13 µV). During the retention interval, the raw EEG signal was digitally bandpass-filtered between 0.1 and 4 Hz (1st-order Butterworth filter, forward only). An SO event was detected when two criteria were met: (i) the filtered signal crossed the individual threshold in the negative direction within 150 ms following a positive-to-negative zero crossing (falling flank), and (ii) the signal subsequently crossed one-third of the individual threshold in the positive direction within 200 ms (rising flank).

### Closed-loop optogenetic inhibition during online-detected slow oscillations

To restrict optogenetic inhibition to NonREM sleep, frontal EEG and EMG signals were bandpass filtered (0.1-40 Hz and 80-300 Hz, respectively; 1st-order Butterworth) and visually monitored together with the video recording to identify NonREM epochs using the same criteria as offline scoring (see *Offline sleep scoring*). Closed-loop SO detection and LED stimulation were enabled only during these periods.

In the In-Phase inhibition condition, a 1 s LED pulse was triggered immediately after an SO was detected. No further SO detections were permitted during the 1 s stimulation window. In the Out-of-Phase-inhibition condition, the LED pulse was triggered after a random delay of 1.5-2.0 s following SO detection. If a new SO was detected before the scheduled pulse onset, the pulse was postponed by 1 s. When the filtered EEG crossed the individual negative threshold within 150 ms following a positive-to-negative zero crossing (falling flank) while the LED was active, the pulse was immediately terminated, and the missed stimulation time was added to subsequent LED pulses (up to a maximum pulse duration of 2 s). If multiple LED pulses were postponed (e.g., during SO trains), rescheduled pulses were separated by random intervals of 200-700 ms to avoid continuous light delivery exceeding 2 s. This closed-loop design ensured that both the number and total duration of LED stimulations were matched across conditions. In the No-Stimulation control condition, SOs were detected online, but the LED remained off. LED power output was kept at 10-20mW at the fiber tip (Optical Power Meter, PM20, Thorlabs) and was delivered bilaterally.

### Histology

After completion of the experiments, animals were deeply anesthetized and transcardially perfused with 4% paraformaldehyde (PFA) in phosphate-buffered saline. Brains were removed, post-fixed in 4% PFA for at least 24 hours, and then sectioned coronally on a vibratome at 50-80 μm thickness. Sections were mounted with an antifade mounting medium containing DAPI (Vectashield). Optic fiber placement above the hippocampus and Jaws expression was verified post-hoc by GFP fluorescence in dorsal CA1 using a light microscope (Leica DMi8).

### Behavioral assessment

Memory performance was quantified by manually scoring object exploration during encoding and retrieval from video recordings analyzed in ANY-maze (Stoelting Europe). Exploration was defined as the rat being within 1 cm of an object with its nose directed toward it and actively sniffing; leaning on the object without sniffing or being farther than 1 cm away was not counted. All videos were scored by the same blinded, experienced experimenter. Memory retrieval in the object-place recognition task was assessed using a discrimination ratio (DR), calculated as:

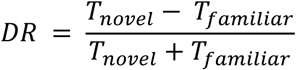

where 𝑇_𝑛𝑜𝑣𝑒𝑙_ and 𝑇_𝑓𝑎𝑚𝑖𝑙𝑖𝑎𝑟_ denote the total exploration time directed toward the object in the novel and familiar location, respectively. A positive discrimination ratio indicates successful memory for the spatial change, whereas a ratio near zero reflects no exploration preference.

### Electrophysiological data analyses

#### Offline sleep scoring

Offline sleep stage classification was performed manually using 10 s epochs, based on one frontal and one parietal EEG channel in combination with the EMG signal. Scoring followed standard criteria ^55^. Wakefulness was characterized by predominant low-amplitude, high-frequency EEG activity accompanied by increased EMG tone. NonREM was defined by high-amplitude delta EEG activity (<4 Hz) and reduced EMG activity. REM sleep was identified by dominant theta EEG activity (4-8 Hz), minimal EMG tone, and the presence of phasic muscle twitches.

#### Offline detection of slow oscillations and sleep spindles

SOs and spindles were detected offline as described previously ^30^. For offline SO detection, EEG signals from NonREM sleep were bandpass-filtered between 0.1-4 Hz (3rd-order Butterworth). SOs were defined by two consecutive positive-to-negative zero crossings separated by 0.5-2.0 s. From all detected events, the 33% with the largest negative peak amplitudes were selected. Spindles were detected after filtering between 10 and 16 Hz (6th-order Butterworth). The smoothed absolute Hilbert transform of the signal was used to identify events exceeding (i) 1.5 SD of the mean NonREM level for 0.5-2.5 s, (ii) 2 SD for 0.25-2.5 s, and (iii) at least once 2.5 SD within the same event.

#### SO-spindle events

The co-occurrence of online-detected SOs and offline-detected sleep spindles was quantified by calculating the rate of spindle onsets occurring within 1 s after an online SO detection, allowing the distinction between solitary and coupled SO-spindle events. To characterize SO-spindle phase-amplitude coupling, the EEG signal was bandpass filtered in the narrow SO range (0.95-1.05 Hz; 3rd-order Butterworth filter), and the instantaneous phase was extracted from the Hilbert-transformed signal. Spindle peaks were identified from the absolute value of the Hilbert-transformed EEG signal, bandpass-filtered in the spindle band, for each offline-detected spindle. If the maximum amplitude of a given spindle occurred within a ±1 s window around an online SO detection, the corresponding SO phase was extracted and stored for further analysis.

#### Spectral analyses

To estimate spectral power across sleep stages, the continuous EEG signal was segmented into 4 s epochs and power spectra were computed over 1-45 Hz with a frequency resolution of 0.05 Hz using fast Fourier transform with a Hanning taper in FieldTrip ^56^. For time-frequency analyses, ±5 s segments centered on online-detected SOs, or on the inhibition onset in the delayed-inhibition condition, were analyzed using a multitaper convolution approach with a Hanning taper. Frequency-specific time windows of seven cycles per frequency were used to estimate power from 5 to 40 Hz in steps of 0.5 Hz. Power values were normalized on a per-animal basis to the average power in a baseline window from −1.5 to −0.5 s relative to SO detection, or from −1 to 0 s relative to inhibition onset in the Out-of-Phase condition.

#### Spike sorting

Spike sorting was performed on acute hippocampal recordings obtained under urethane anesthesia, using SpikeInterface ^57^. Raw extracellular recordings were bandpass filtered between 300 and 6000 Hz, globally re-referenced, and whitened prior to spike detection. Automated spike sorting was carried out using MountainSort5 with default parameters. Resulting units were subjected to automated quality-based curation, and units were retained only if they met all of the following criteria: signal-to-noise ratio >5, inter-spike interval violation ratio <0.2, and nearest-neighbor isolation metric >0.7. All retained units were subsequently visually inspected using the SpikeInterface graphical user interface, and units exhibiting non-physiological waveforms, unstable firing rates, or clear contamination by noise were excluded.

#### Validation of optogenetic inhibition

To quantify optogenetic inhibition under anesthesia, spikes from all recorded single units (N = 116; 3 animals) were counted across inhibition trials (N = 1469) within the inhibition window and a matched baseline window (0.5-1 s). Suppression was calculated as the normalized difference between inhibition and baseline spike counts (−1 = strong suppression; 0 = no change; >0 = increased firing). For statistical analysis, firing rates were averaged across trials per unit and compared between windows using a Wilcoxon signed-rank test.

### Statistical analyses

Statistical analyses were performed using custom scripts in R ^58^ or MATLAB (Version 2023b). Animals were excluded a priori for (1) insufficient Jaws-GFP expression, (2) incorrect optic fiber placement above CA1, or (3) low encoding exploration (<1 s per object). Based on these criteria, three animals were excluded for insufficient expression and one for enlarged ventricles. Statistical comparisons across conditions were conducted using linear mixed models, fitted using the lme4 package ^59^, with rat as a random intercept and group factors as fixed effects. To compare memory performance, for instance, inhibition condition (in-phase vs. out-of-phase vs. no-stimulation) and condition order (whether an inhibition condition was tested first, second, or third in the within design) were compared using the formula:

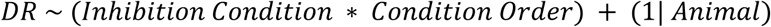

where DR indicated the discrimination ratio over the 5 min test interval. The significance of factors was assessed by stepwise removal of the respective main effect or interaction from the model and comparison of nested models using likelihood-ratio tests. Behavioral control parameters - including total distance traveled, sleep duration, and total object exploration time during encoding and retrieval phases - were analyzed analogously. Post-hoc comparisons were performed using two-sided Welch’s t-tests. Correlations were assessed using Spearman’s rank coefficients to account for the small sample size.

SO-spindle phase-amplitude coupling was assessed using Rayleigh tests for non-uniformity of circular distributions, as implemented in the Circular Statistics Toolbox ^60^, and group differences were evaluated using Watson-Williams tests as implemented in the circular package ^61^. Time-frequency representations were compared across conditions and against baseline using dependent-samples *t*-tests with cluster-based permutation correction (Monte Carlo method, 5,000 permutations, two-sided) as implemented in FieldTrip ^56^. Mediation analysis was performed by estimating indirect and direct effects using quasi-Bayesian Monte Carlo simulation as implemented in the mediation framework ^62^. Linear mixed-effects models with animal as a random intercept were fitted for the total, mediator, and outcome paths, and mediation effects were estimated from 1,000 simulations. For all analyses a *p* < 0.05 was considered significant. Results were visualized using ggplot2 ^63^ and ggpubr ^64^ packages.

## Data availability

All data and code required to reproduce the statistical analyses and plots in the paper are available at the following repository: https://github.com/MaxHarkotte/Hippocampus-consolidates-memory-in-the-upstate-of-cortical-sleep-slow-oscillations. Any further materials will be made available upon reasonable request.

## Author contributions

M.H., M.I., J.B. and N.N. designed the study. M.H., J.B. and N.N. wrote the paper. M.H. collected and analyzed the data.

## Acknowledgments

We thank Francesco Gobbo for sharing protocols for the use of Jaws in rats. We are grateful to Edward S. Boyden for making the viral construct available. We also thank Ilona Sauter, Daniel Gramling and Klaus Vollmer for technical support.

## Funding

This study was supported by grants from the Deutsche Forschungsgemeinschaft to J.B. (FOR 5434) and the European Research Council to J.B. (ERC AdG 883098 SleepBalance). M.I. and N.N. are supported by the Hertie Foundation (Hertie Network of Excellence in Clinical Neuroscience).

## Extended Data Figures

**Extended Data Fig. 1.**
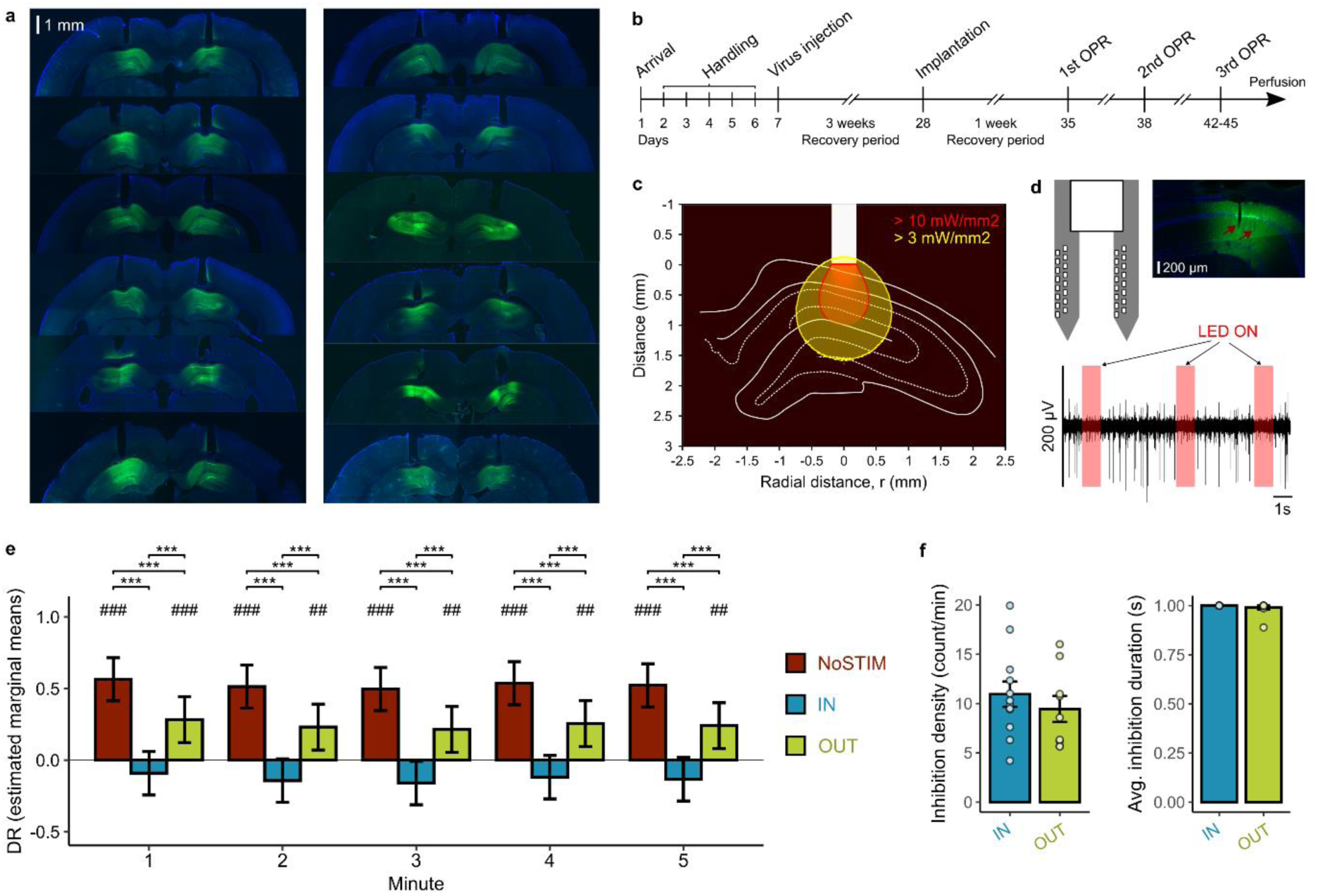
Validation of hippocampal inhibition and memory performance controlling for testing order. **a,** *Histology*. Bilateral optic fibers positioned above dorsal CA1 in animals expressing the red-light–activated chloride pump Jaws-GFP (n = 12). **b,** *Experimental timeline.* Animals were handled for 5 days prior to the first surgery, during which Jaws-GFP was injected bilaterally into the hippocampus. After 3 weeks of recovery, a second surgery was performed to implant EEG and EMG electrodes together with bilateral optic fibers above the dorsal hippocampus. Following a 1-week recovery period, animals were repeatedly tested in the object-recognition task; the first two tests were separated by 2 days and the third by 3-7 days to reduce interference. Animals were subsequently transcardially perfused. **c,** *Light delivery.* Estimated light spread within the hippocampus from a 625-nm LED pulse delivered through a 400-µm optic fiber. Light propagation was modeled using a previously published MATLAB toolbox ^65^. **d,** *Acute recordings.* Top left: A two-shank, 32-channel sharpened silicon probe (P2-ASSY-116, Cambridge Neurotech; grey) mounted to a 400-µm optic fiber (white) was used during acute recordings. Top right: Histological verification of probe shank placement in CA1 expressing Jaws-GFP following acute recording. Bottom, representative bandpass-filtered trace (300–3000 Hz, 4th-order Butterworth) showing three trials of hippocampal inhibition (red). **e,** *Memory performance corrected for testing order.* To confirm whether the effect of closed-loop optogenetic inhibition locked to online-detected SOs (No-Stimulation vs. In-Phase vs. Out-of-Phase) did not depend on the order in which the conditions were tested, estimated marginal means were derived from a linear mixed-effects model including testing order as a factor (in addition to inhibition condition as well as minute across the retrieval phase as fixed factors and animal as random intercept). While a significant effect of testing order was found (χ²(2) = 8.84, *p* = 0.012), the main effect of inhibition condition on memory performance remained significant (χ²(2) = 97.45, *p* < 0.001). No effect was found for the minute during the retrieval phase, from which the discrimination ratio (DR) was calculated (χ²(4) = 0.93, *p* = 0.92), although minute was retained in the model for completeness. Bars indicate the estimated marginal means across all five minutes of the retrieval phase, adjusting for testing order, and error bars represent SEM. Symbols above bars denote model-based tests of estimated marginal means against zero (^##^p< 0.01, ^###^p< 0.001); brackets indicate pairwise condition contrasts (***p< 0.001). **f,** *Inhibition across conditions.* Inhibition density (left) and average duration of one inhibition (right) were comparable between the In-Phase and Out-of-Phase conditions.

**Extended Data Fig. 2.**
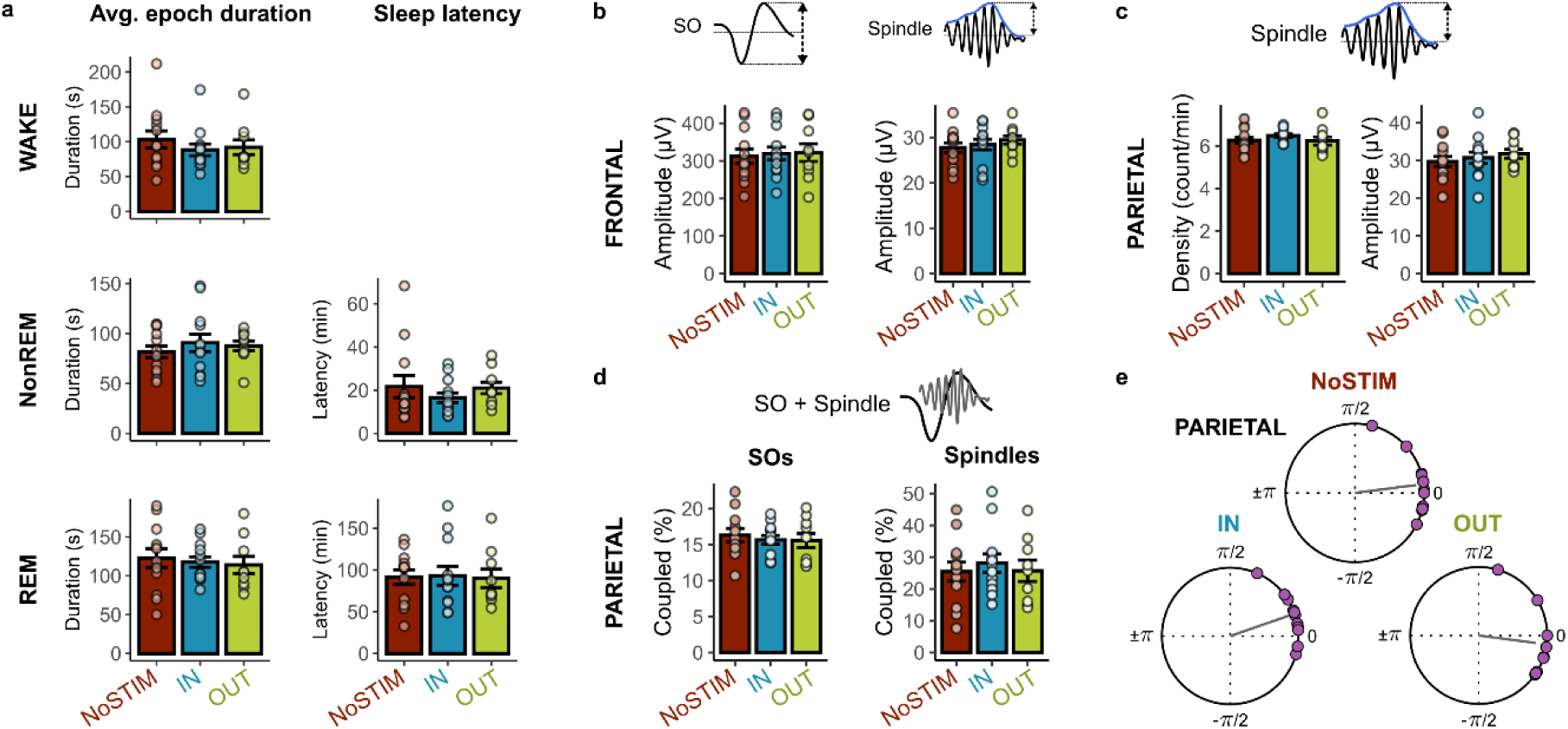
Control measures of macro sleep architecture and sleep-oscillatory activity show no difference despite hippocampal closed-loop inhibition. **a,** Left, Average epoch duration of Wake, NonREM, and REM sleep. Right, Sleep latency to NonREM, and REM sleep. All measures were comparable across conditions (all *p* > 0.05). **b,** Negative-to-positive peak amplitude of online detected SOs (left) and absolute peak amplitude of sleep spindles (right) detected on the left frontal EEG electrode were comparable across conditions. **c,** Spindle density and peak amplitudes were comparable across conditions also when detected in parietal EEG, i.e., closer to the hippocampal inhibition. **d,** SO-spindle coupling was comparable across conditions in the parietal EEG. **e,** Mean SO phase at which spindle peak powered in parietal EEG. Purple dots indicate individual animals; grey lines indicate the mean phase of maximal spindle power. In all conditions, spindle power peaked near the SO positive maximum (0°, Watson-Williams Test for condition comparison: *p* = 0.13).

**Extended Data Fig. 3.**
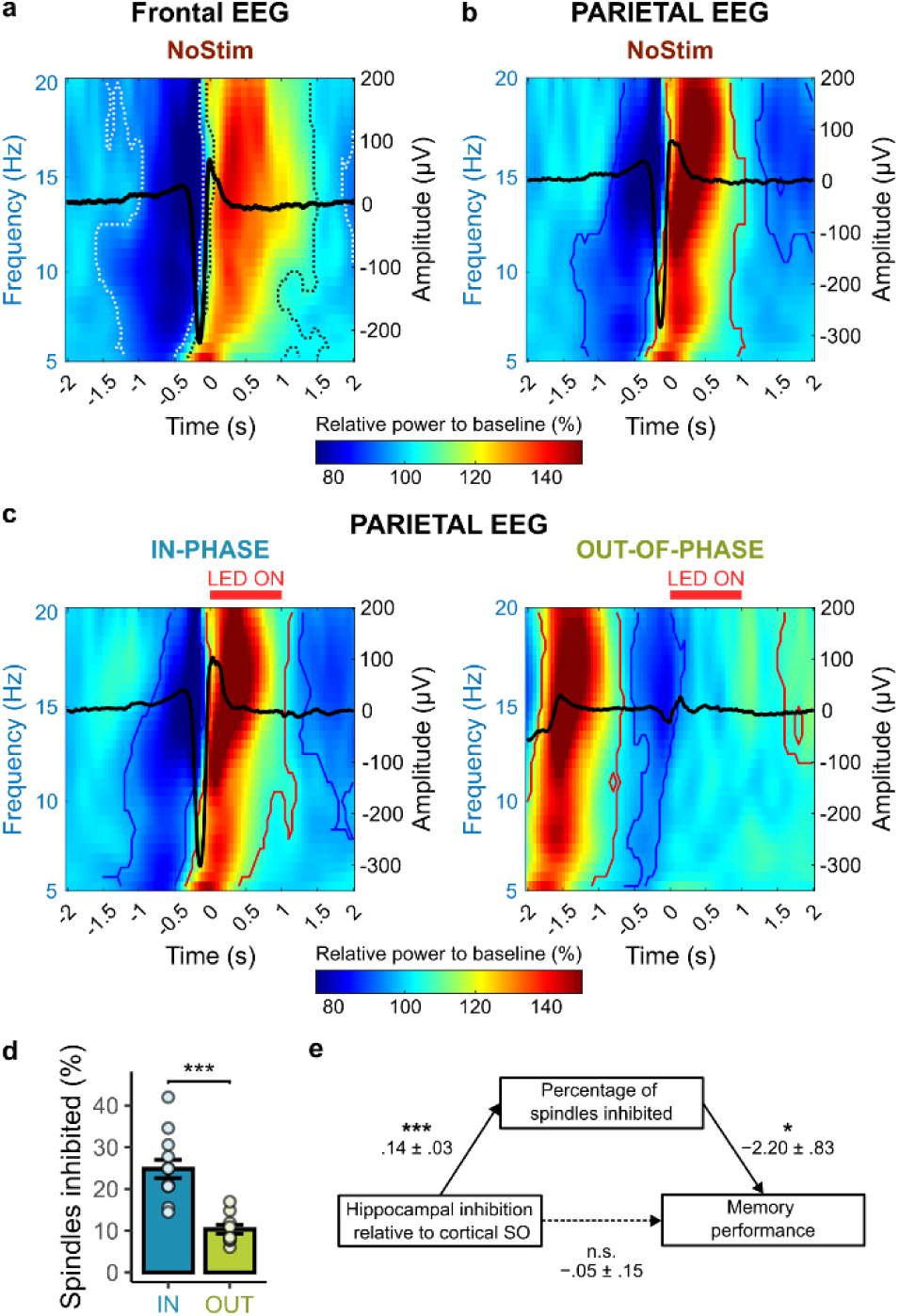
Spindle-band power increase during the SO upstate under undisturbed conditions and spindle-mediated memory impairment in parietal EEG. **a,** Time-frequency representation locked to the online-detection of a SO in the No-Stimulation control condition for frontal EEG. Dashed lines indicate significant positive (black) and negative (white) spectral clusters relative to baseline. **b,** Same as (a), but for parietal EEG. **c,** Same as (a), but time-frequency representations are locked to the inhibition onset in the In-Phase (left) and Out-of-Phase (right) conditions for parietal EEG. **d,** The fraction of all parietal spindles overlapping with the hippocampal inhibition window was significantly higher in the In-Phase than in the Out-of-Phase condition (*t*(16.20) = 5.58, \*\*\**p* < 0.001). **e,** Mediation model relating inhibition condition (SO in-phase versus Out-of-Phase), the percentage of inhibited spindles in parietal EEG, and memory performance. Path coefficients represent unstandardized fixed-effect estimates (estimate ± s.e.) from linear mixed-effects models with animals included as a random intercept. Solid arrows indicate significant paths; the dashed arrow denotes the direct effect of condition after accounting for the mediator (*p< 0.05, ***p< 0.001).

## Notes

### Competing Interest Statement

The authors have declared no competing interest.

## References

1. Stickgold, R. Sleep-dependent memory consolidation. Nature 437, 1272–1278 (2005).

2. Lutz, N. D., Harkotte, M. & Born, J. Sleep’s contribution to memory formation. Physiological Reviews 10.1152/physrev.00054.2024 (2025) doi:10.1152/physrev.00054.2024.

3. Buzsáki, G. The Hippocampo-Neocortical Dialogue. Cereb Cortex 6, 81–92 (1996).

4. Diekelmann, S. & Born, J. The memory function of sleep. Nat Rev Neurosci 11, 114–126 (2010).

5. Klinzing, J. G., Niethard, N. & Born, J. Mechanisms of systems memory consolidation during sleep. Nature Neuroscience 22, 1598–1610 (2019).

6. Staresina, B. P., Niediek, J., Borger, V., Surges, R. & Mormann, F. How coupled slow oscillations, spindles and ripples coordinate neuronal processing and communication during human sleep. Nat Neurosci 26, 1429–1437 (2023).

7. Rothschild, G., Eban, E. & Frank, L. M. A cortical–hippocampal–cortical loop of information processing during memory consolidation. Nat Neurosci 20, 251–259 (2017).

8. Brodt, S., Inostroza, M., Niethard, N. & Born, J. Sleep-A brain-state serving systems memory consolidation. Neuron 111, 1050–1075 (2023).

9. Puentes-Mestril, C., Roach, J., Niethard, N., Zochowski, M. & Aton, S. J. How rhythms of the sleeping brain tune memory and synaptic plasticity. Sleep 42, zsz095 (2019).

10. Massimini, M., Huber, R., Ferrarelli, F., Hill, S. & Tononi, G. The Sleep Slow Oscillation as a Traveling Wave. J. Neurosci. 24, 6862–6870 (2004).

11. Steriade, M. Grouping of brain rhythms in corticothalamic systems. Neuroscience 137, 1087–1106 (2006).

12. Marshall, L., Helgadóttir, H., Mölle, M. & Born, J. Boosting slow oscillations during sleep potentiates memory. Nature 444, 610–613 (2006).

13. Miyamoto, D. Optical imaging and manipulation of sleeping-brain dynamics in memory processing. Neuroscience Research 181, 9–16 (2022).

14. Steriade, M., Timofeev, I. & Grenier, F. Natural Waking and Sleep States: A View From Inside Neocortical Neurons. Journal of Neurophysiology 85, 1969–1985 (2001).

15. Volgushev, M., Chauvette, S., Mukovski, M. & Timofeev, I. Precise Long-Range Synchronization of Activity and Silence in Neocortical Neurons during Slow-Wave Sleep. J. Neurosci. 26, 5665–5672 (2006).

16. Sirota, A., Csicsvari, J., Buhl, D. & Buzsáki, G. Communication between neocortex and hippocampus during sleep in rodents. Proceedings of the National Academy of Sciences 100, 2065–2069 (2003).

17. Helfrich, R. F. et al. Bidirectional prefrontal-hippocampal dynamics organize information transfer during sleep in humans. Nat Commun 10, 3572 (2019).

18. Niknazar, H., Malerba, P. & Mednick, S. C. Slow oscillations promote long-range effective communication: The key for memory consolidation in a broken-down network. Proceedings of the National Academy of Sciences 119, e2122515119 (2022).

19. Wilson, M. A. & McNaughton, B. L. Reactivation of Hippocampal Ensemble Memories During Sleep. Science 265, 676–679 (1994).

20. Buzsáki, G. Hippocampal sharp wave-ripple: A cognitive biomarker for episodic memory and planning. Hippocampus 25, 1073–1188 (2015).

21. Schreiner, T. & Staudigl, T. Electrophysiological signatures of memory reactivation in humans. Philos Trans R Soc Lond B Biol Sci 375, 20190293 (2020).

22. Robinson, H. L. et al. Large sharp-wave ripples promote hippocampo-cortical memory reactivation and consolidation during sleep. Neuron 114, 226–236.e6 (2026).

23. Staresina, B. P. et al. Hierarchical nesting of slow oscillations, spindles and ripples in the human hippocampus during sleep. Nat Neurosci 18, 1679–1686 (2015).

24. Klinzing, J. G. et al. Spindle activity phase-locked to sleep slow oscillations. NeuroImage 134, 607–616 (2016).

25. Clemens, Z. et al. Temporal coupling of parahippocampal ripples, sleep spindles and slow oscillations in humans. Brain 130, 2868–2878 (2007).

26. Fernandez, L. M. J. & Lüthi, A. Sleep Spindles: Mechanisms and Functions. Physiological Reviews 100, 805–868 (2020).

27. Peyrache, A. & Seibt, J. A mechanism for learning with sleep spindles. Philos Trans R Soc Lond B Biol Sci 375, 20190230 (2020).

28. Maingret, N., Girardeau, G., Todorova, R., Goutierre, M. & Zugaro, M. Hippocampo-cortical coupling mediates memory consolidation during sleep. Nature Neuroscience 19, 959–964 (2016).

29. Binder, S. et al. Sleep enhances memory consolidation in the hippocampus-dependent object-place recognition task in rats. Neurobiology of Learning and Memory 97, 213–219 (2012).

30. Sawangjit, A. et al. The hippocampus is crucial for forming non-hippocampal long-term memory during sleep. Nature 564, 109–113 (2018).

31. Oyanedel, C. N. et al. Role of slow oscillatory activity and slow wave sleep in consolidation of episodic-like memory in rats. Behav Brain Res 275, 126–130 (2014).

32. Bolsius, Y. G. et al. Recovering object-location memories after sleep deprivation-induced amnesia. Current Biology 33, 298–308.e5 (2023).

33. Takashima, A. et al. Declarative memory consolidation in humans: A prospective functional magnetic resonance imaging study. Proceedings of the National Academy of Sciences 103, 756–761 (2006).

34. Sawangjit, A., Oyanedel, C. N., Niethard, N., Born, J. & Inostroza, M. Deepened sleep makes hippocampal spatial memory more persistent. Neurobiology of Learning and Memory 173, 107245 (2020).

35. Wilhelm, I. et al. Sleep selectively enhances memory expected to be of future relevance. J Neurosci 31, 1563–1569 (2011).

36. Oyanedel, C. N., Durán, E., Niethard, N., Inostroza, M. & Born, J. Temporal associations between sleep slow oscillations, spindles and ripples. Eur J Neurosci 52, 4762–4778 (2020).

37. Sanda, P. et al. Bidirectional Interaction of Hippocampal Ripples and Cortical Slow Waves Leads to Coordinated Spiking Activity During NREM Sleep. Cereb Cortex 31, 324–340 (2021).

38. Peyrache, A., Khamassi, M., Benchenane, K., Wiener, S. I. & Battaglia, F. P. Replay of rule-learning related neural patterns in the prefrontal cortex during sleep. Nat Neurosci 12, 919–926 (2009).

39. Latchoumane, C.-F. V., Ngo, H.-V. V., Born, J. & Shin, H.-S. Thalamic Spindles Promote Memory Formation during Sleep through Triple Phase-Locking of Cortical, Thalamic, and Hippocampal Rhythms. Neuron 95, 424–435.e6 (2017).

40. Ngo, H.-V. V., Martinetz, T., Born, J. & Mölle, M. Auditory Closed-Loop Stimulation of the Sleep Slow Oscillation Enhances Memory. Neuron 78, 545–553 (2013).

41. Kim, J., Gulati, T. & Ganguly, K. Competing Roles of Slow Oscillations and Delta Waves in Memory Consolidation versus Forgetting. Cell 179, 514–526.e13 (2019).

42. Girardeau, G., Benchenane, K., Wiener, S. I., Buzsáki, G. & Zugaro, M. B. Selective suppression of hippocampal ripples impairs spatial memory. Nat Neurosci 12, 1222–1223 (2009).

43. Ego-Stengel, V. & Wilson, M. A. Disruption of ripple-associated hippocampal activity during rest impairs spatial learning in the rat. Hippocampus 20, 1–10 (2010).

44. Gridchyn, I., Schoenenberger, P., O’Neill, J. & Csicsvari, J. Assembly-Specific Disruption of Hippocampal Replay Leads to Selective Memory Deficit. Neuron 106, 291–300.e6 (2020).

45. Hahn, T. T. G., Sakmann, B. & Mehta, M. R. Phase-locking of hippocampal interneurons’ membrane potential to neocortical up-down states. Nat Neurosci 9, 1359–1361 (2006).

46. Hahn, T. T. G., Sakmann, B. & Mehta, M. R. Differential responses of hippocampal subfields to cortical up–down states. Proceedings of the National Academy of Sciences 104, 5169–5174 (2007).

47. Isomura, Y. et al. Integration and Segregation of Activity in Entorhinal-Hippocampal Subregions by Neocortical Slow Oscillations. Neuron 52, 871–882 (2006).

48. Swanson, R. A. et al. Topography of putative bi-directional interaction between hippocampal sharp-wave ripples and neocortical slow oscillations. Neuron 113, 754–768.e9 (2025).

49. Niethard, N., Ngo, H.-V. V., Ehrlich, I. & Born, J. Cortical circuit activity underlying sleep slow oscillations and spindles. PNAS 115, E9220–E9229 (2018).

50. Siapas, A. G. & Wilson, M. A. Coordinated interactions between hippocampal ripples and cortical spindles during slow-wave sleep. Neuron 21, 1123–1128 (1998).

51. Mölle, M., Eschenko, O., Gais, S., Sara, S. J. & Born, J. The influence of learning on sleep slow oscillations and associated spindles and ripples in humans and rats. Eur J Neurosci 29, 1071–1081 (2009).

52. Karaba, L. A. et al. A hippocampal circuit mechanism to balance memory reactivation during sleep. Science 385, 738–743 (2024).

53. Vyazovskiy, V. V., Riedner, B. A., Cirelli, C. & Tononi, G. Sleep Homeostasis and Cortical Synchronization: II. A Local Field Potential Study of Sleep Slow Waves in the Rat. Sleep 30, 1631–1642 (2007).

54. Nir, Y. et al. Regional Slow Waves and Spindles in Human Sleep. Neuron 70, 153–169 (2011).

55. Neckelmann, D., Olsen, Ø. E., Fagerland, S. & Ursin, R. The Reliability and Functional Validity of Visual and Semiautomatic Sleep/Wake Scoring in the Møll-Wistar Rat. Sleep 17, 120–131 (1994).

56. Oostenveld, R., Fries, P., Maris, E. & Schoffelen, J.-M. FieldTrip: Open Source Software for Advanced Analysis of MEG, EEG, and Invasive Electrophysiological Data. Comput Intell Neurosci 2011, 156869 (2011).

57. Buccino, A. P. et al. SpikeInterface, a unified framework for spike sorting. eLife 9, e61834 (2020).

58. R Core Team. R: A Language and Environment for Statistical Computing. R Foundation for Statistical Computing (2024).

59. Bates, D., Mächler, M., Bolker, B. & Walker, S. Fitting Linear Mixed-Effects Models Using lme4. Journal of Statistical Software 67, 1–48 (2015).

60. Berens, P. CircStat: A MATLAB Toolbox for Circular Statistics. Journal of Statistical Software 31, 1–21 (2009).

61. Agostinelli, C. & Lund, U. R package ‘circular’: Circular Statistics. (2025).

62. Tingley, D., Yamamoto, T., Hirose, K., Keele, L. & Imai, K. mediation: R Package for Causal Mediation Analysis. Journal of Statistical Software 59, 1–38 (2014).

63. Wickham, H. ggplot2. WIREs Computational Statistics 3, 180–185 (2011).

64. Kassambara, A. ggpubr: ‘ggplot2’ Based Publication Ready Plots. (2025).

65. Stujenske, J. M., Spellman, T. & Gordon, J. A. Modeling the Spatiotemporal Dynamics of Light and Heat Propagation for In Vivo Optogenetics. Cell Reports 12, 525–534 (2015).

